# Touch to text: Spatiotemporal evolution of braille letter representations in blind readers

**DOI:** 10.1101/2024.10.30.620429

**Authors:** Santani Teng, Radoslaw Cichy, Dimitrios Pantazis, Aude Oliva

## Abstract

Visual deprivation does not silence the visual cortex, which is responsive to auditory, tactile, and other nonvisual tasks in blind persons. However, the underlying functional dynamics of the neural networks mediating such crossmodal responses remain unclear. Here, using braille reading as a model framework to investigate these networks, we presented sighted (N=13) and blind (N=12) readers with individual visual print and tactile braille alphabetic letters, respectively, during MEG recording. Using time-resolved multivariate pattern analysis and representational similarity analysis, we traced the alphabetic letter processing cascade in both groups of participants. We found that letter representations unfolded more slowly in blind than in sighted brains, with decoding peak latencies ∼200 ms later in braille readers. Focusing on the blind group, we found that the format of neural letter representations transformed within the first 500 ms after stimulus onset from a low-level structure consistent with peripheral nerve afferent coding to high-level format reflecting pairwise letter embeddings in a text corpus. The spatiotemporal dynamics of the transformation suggest that the processing cascade proceeds from a starting point in somatosensory cortex to early visual cortex and then to inferotemporal cortex. Together our results give insight into the neural mechanisms underlying braille reading in blind persons and the dynamics of functional reorganization in sensory deprivation.

## INTRODUCTION

In the absence of vision, cortical areas traditionally characterized as visual in sighted people are recruited for a range of nonvisual tasks (Pascual-Leone et al., 2005; Merabet and Pascual-Leone, 2010; Kupers and Ptito, 2014; Amedi et al., 2017; Bedny, 2017), suggesting a functional reorganization in response to sensory deprivation. For decades, such crossmodal plasticity in blindness has been investigated using the cornerstone model of braille, a text system of raised dots designed to be read haptically with the fingerpad (Millar, 1997; Englebretson et al., 2023). This work has revealed a cortical network of braille-sensitive regions including the left-lateralized “Visual” Word Form Area (VWFA) (Dehaene and Cohen, 2011) representing multimodal orthographic formats and visual print (Reich et al., 2011; Striem-Amit et al., 2012; Kim et al., 2017; Rączy et al., 2019), and even early visual cortex (EVC), the starting point of the standard cortical visual processing hierarchy (Sadato et al., 1996; Reich et al., 2011). Notably, EVC activity is not merely epiphenomenal, but functionally important to braille reading: disruption, whether by stroke or magnetic stimulation, selectively impairs letter recognition while sparing basic somatosensation (Cohen et al., 1997; Hamilton and Pascual-Leone, 1998; Hamilton et al., 2000). However, the extent of functional reorganization in, and thus the functional role of, regions like EVC in the brains of congenitally blind persons remains a subject of ongoing debate (Amedi et al., 2017; Bedny, 2017; Makin and Krakauer, 2023; Seydell-Greenwald et al., 2023), in part becausethe temporal and representational dynamics of the braille processing network remain poorly understood.

A larger body of analogous work has revealed how, in the sighted brain, print letters as a special class of visual object stimulus (Cichy et al., 2014; Bi et al., 2016) are represented as patterns of retinal stimulation, sub-letter features, and distinct and abstract letter identities in high-level ventral visual cortex (Grainger et al., 2008; Madec et al., 2012; Thesen et al., 2012; Isik et al., 2014; Fischer-Baum et al., 2017). Similarly elucidating the dynamics of the braille text perception network in blind readers would clarify both the building blocks of the braille reading process as well as structure of the larger sensory processing hierarchy in a functionally reorganized brain. This requires describing not only the anatomical loci of relevant brain activity <u>(Reich et al. 2011; Liu, Rapp, and</u> <u>Bedny 2023)</u>, but the temporal and representational dynamics of the process. In analogy to print characters, our working hypothesis was that braille characters must likewise develop in ordered fashion from dot patterns impinging on the fingerpad into meaningful components of written language, mediated by a cortical network including EVC (Hamilton and Pascual-Leone, 1998; Ioannides et al., 2013; Haupt et al., 2024).

To test this hypothesis and reveal the underlying neural networks, we presented blind participants with braille letters and sighted participants with visual print letters while recording brain responses with MEG. We then used multivariate pattern classification (Haynes and Rees, 2006; Carlson et al., 2013; Cichy et al., 2014) and model comparison using representational similarity analysis (RSA) (Kriegeskorte et al., 2008; Kriegeskorte and Kievit, 2013) to low- and high-level models of alphabetic letter representation to characterize the processing hierarchy in both sighted and blind readers.

## RESULTS

We presented congenitally or early-blind (N=12) and typically sighted (N=13) participants with lowercase alphabetic letters while recording brain activity with magnetoencephalography (MEG). We used analogous experimental designs and analyses of each group: both subject groups were presented with a common set of 10 different consonants, in tactile braille and in visual printed formats respectively (Fig. 1a; purple eye and green hand icons indicate procedures and results for Sighted and Blind participant groups, respectively, hereafter). Blind participants were all braille-proficient since early childhood and had no history of vision better than nonspatial light perception, and thus had no visual experience of the printed alphabet (see Table 1 for blind participant details). They were presented with braille letters via a refreshable tactile braille cell to the stationary fingerpad of their preferred index finger (10 left, 2 right). Sighted subjects had normal or corrected-to-normal vision, no experience with reading braille visually, and were presented with lowercase print letters in standard font centrally on a computer screen rear-projected into the MEG chamber.

**Figure 1.**
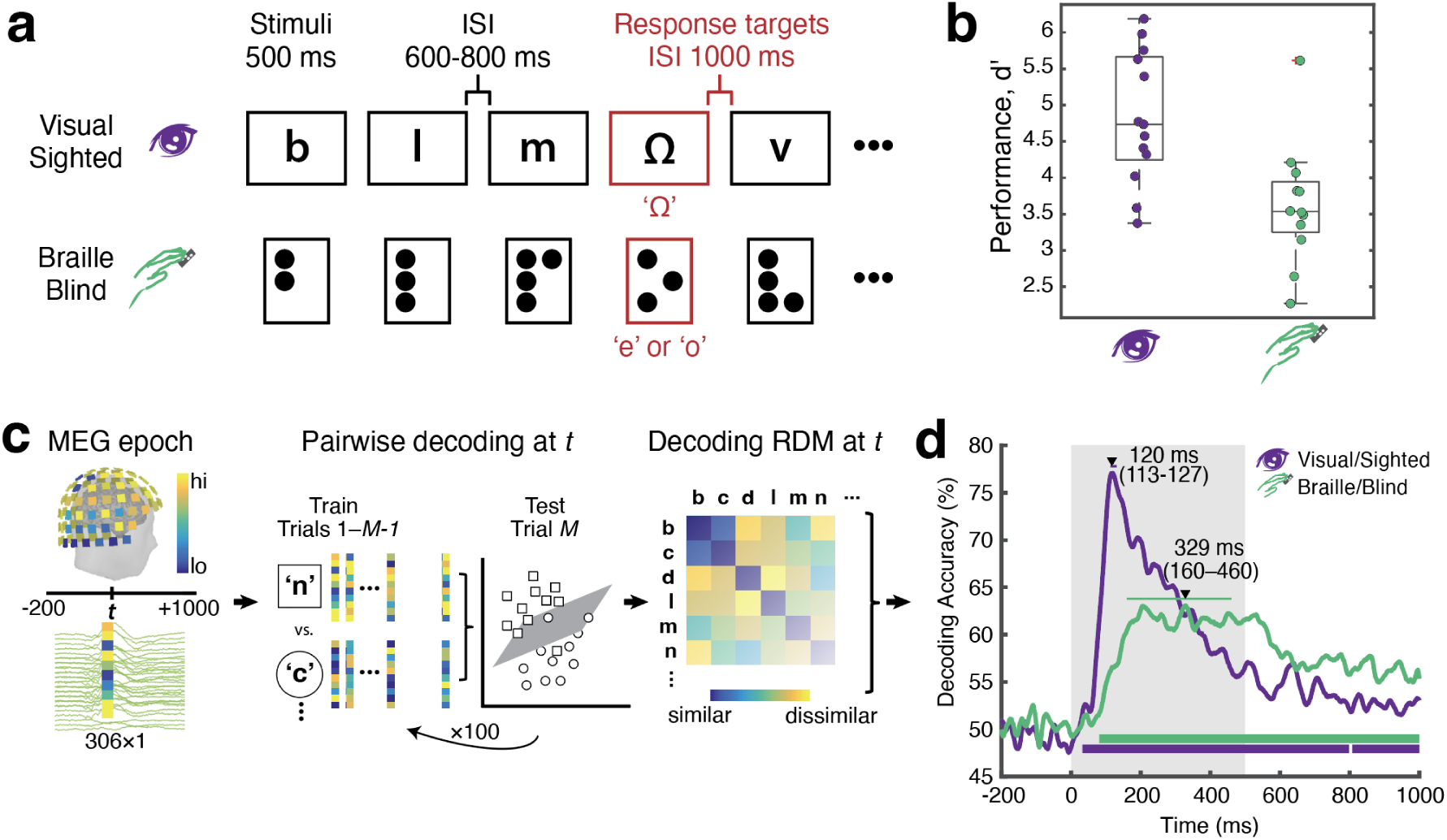
Experimental setup, analysis, and general MEG decoding results. (**a**) Stimulus presentation and trial schematic. Single alphabetical letters were presented visually at central fixation to sighted subjects, or to the stationary index fingerpad via braille display to blind subjects. Procedures and results for the Sighted and Blind groups are color-coded and denoted with purple eye and green hand icons, respectively. (**b**) Both groups readily perceived and responded to occasional vigilance targets. Color-coded scatter plots indicate *d*’ for individuals; boxplot shows median and interquartile range. (**c**) Time-resolved multivariate pattern analysis. For each time point *t* and trial separately, we arranged MEG activity across the 306 sensors into MEG pattern vectors. Single-trial pattern vectors were then whitened, and randomly assigned to bins that were averaged to create *M* pseudo-trial vectors for increased signal-to-noise ratio. We classified experimental conditions (i.e. the 10 letters) pairwise from the pseudotrials using a linear support vector machine (SVM) classifier. The assignment of raw trials to pseudotrials and classification procedure was repeated 100 times; the resulting mean decoding accuracy for each stimulus pair populated a decoding matrix indexed in rows and conditions by the letters classified, for each *t*. The decoding matrix — a representational dissimilarity matrix — formed the basis of further analyses. For example, averaging across the matrix yielded a grand average time course indicating the dynamics with which letter representations emerge. (**d**) Grand-average classification time courses for letters for blind (green) and sighted (purple) participants. Color-coded bars beneath plots indicate clusters of significant decoding; arrows indicate peaks, with thin horizontal bars indicating 95% confidence intervals (CIs). Shaded region from 0–500 ms indicates stimulus duration period. Significance was assessed via permutation-based cluster-size inference (p<0.05 cluster-definition threshold, p<0.05 cluster threshold, one-sided, 500 permutations).

**Table 1.** Blind participant demographics and characteristics. *Total* = total blindness, no light perception in either eye. *LP* = light perception, no spatial perception.

| ID | M/F | Age at test (years) | Blindness level | Blindness onset age (years) |
| --- | --- | --- | --- | --- |
| 1 | F | 34 | Total | 0 |
| 2 | F | 24 | LP | 0 |
| 3 | M | 28 | Total | 0 |
| 4 | F | 29 | Total | 0 |
| 5 | M | 27 | LP | 0 |
| 6 | F | 26 | LP | 0 |
| 7 | F | 29 | Total | 0 |
| 8 | F | 32 | Total | 3 |
| 9 | M | 39 | LP | 0 |
| 10 | M | 24 | LP | 0 |
| 11 | F | 35 | LP | 3 |
| 12 | M | 18 | LP | <1 |

On each trial, participants were presented with a letter for 500ms, followed by a 600 – 800 ms interstimulus interval (ISI). Trials were presented in random order, interspersed with vigilance trials on which participants responded to a target character (either an ‘e’ or an ‘o’ for blind, and ‘omega’ (Ω) for sighted) with a button press. Vigilance trials were excluded from further analysis.

Both blind and sighted participants identified letters easily, as indicated by informal pretesting and high performance on the vigilance task (mean hit rate 90%, *d*’ 3.63 for blind; hit rate 96%, *d*’ 4.83 for sighted; see Fig. 1b). Notably, blind participants performed highly despite receiving the stimulus on a static finger, rather than making sweeping motions typical of naturalistic braille text reading. This allowed us to precisely mark the experimenter-defined stimulus onset in both blind and sighted participants.

### The temporal dynamics of braille and visual letter identity

To determine the temporal dynamics with which letter representations emerge in sighted and blind brains, we subjected both data sets to an analogous analysis pipeline (Fig. 1c). We first extracted trial epochs spanning -200 ms to +1000 ms relative to stimulus onset. We then used time-resolved multivariate pattern analysis (MVPA) (Carlson et al., 2013; Cichy et al., 2014; Guggenmos et al., 2018) to classify the 10 letter conditions pairwise from MEG data for every millisecond in the epoch. We saved the classification results (i.e. decoding accuracy) in a 10 x 10 decoding accuracy matrix (symmetric across the diagonal, and the diagonal undefined) indexed in rows and columns by the conditions classified.

Averaging across decoding accuracy matrices at each point, we obtained grand-average time courses indicating the dynamics with which letter representations emerge in blind and sighted subjects (Fig. 1c, *right* panel). We assessed significance using non-parametric sign-permutation tests and controlled for multiple comparisons by cluster correction (Maris and Oostenveld, 2007) (cluster definition threshold p < 0.05, cluster threshold p < 0.05). We report peak latencies as a measure of the time at which representations are best discriminated indicating untangling (DiCarlo and Cox, 2007), as well as onset latencies of the first significant cluster indicating the beginning of distinct letter processing (both with bootstrapped 95% confidence intervals in brackets).

We found that letter stimuli were significantly discriminated in both blind braille readers and sighted visual readers (Fig. 1d). As expected, in the sighted group, we observed the standard curve shape for classifying single visual stimuli with an early onset at 32 ms (22–75 ms), precipitating a sharp rise to a peak at 120 ms (113–127 ms), followed by a gradual decline. In contrast, in the blind group we observed a different shape of the classification time course: a less steep rise with a comparably early onset at 82 ms (50–125 ms), led to a peak within a wide plateau at 329 ms (160–460 ms). The qualitative differences and similarities were ascertained statistically (Supplementary Table S1), with a significantly later peak by 209 ms (54–390 ms) in the blind compared to the sighted subjects (p < 0.002, determined by bootstrapping) and no significant difference in onset latency (p = 0.08).

Together, these results establish the feasibility of assessing braille letter representations from MEG data, determine the time course with which braille letter representations emerge in the blind brain, and provide a first comparative characterization of the neural dynamics of braille letter representations in relation to visual letter representations in the sighted brain.

### The representational format of neural responses to visual and braille letters

The rationale in using RSA to relate the models to human brain activity (Fig. 2b) is to abstract from incommensurate signal spaces, such as MEG sensors and computational models, to a common space defined by pairwise dissimilarities between experimental conditions. The pairwise dissimilarities are aggregated in *representational dissimilarity matrices* (RDMs) indexed in rows and columns by the conditions compared. Interpreting decoding accuracy as a dissimilarity measure (i.e., higher accuracy indexes greater representational differences (Kriegeskorte et al., 2008)), we used the decoding accuracy matrices from the MVPA analysis as MEG RDMs. To compute the *visual* and *tactile* model RDMs, we presented the experimental stimuli to the models, extracted the resulting activation values into pattern vectors, and calculated pairwise dissimilarity between pattern vectors using 1–Spearman correlation distance. The *bigram* model RDM was populated by the inverted average co-occurrence frequency of each letter pair. We then related the time-resolved MEG RDMs to the model RDMs by computing semipartial Spearman correlations between them, resulting in two correlation time courses for each participant group (Fig. 2b, *right*).

**Figure 2:**
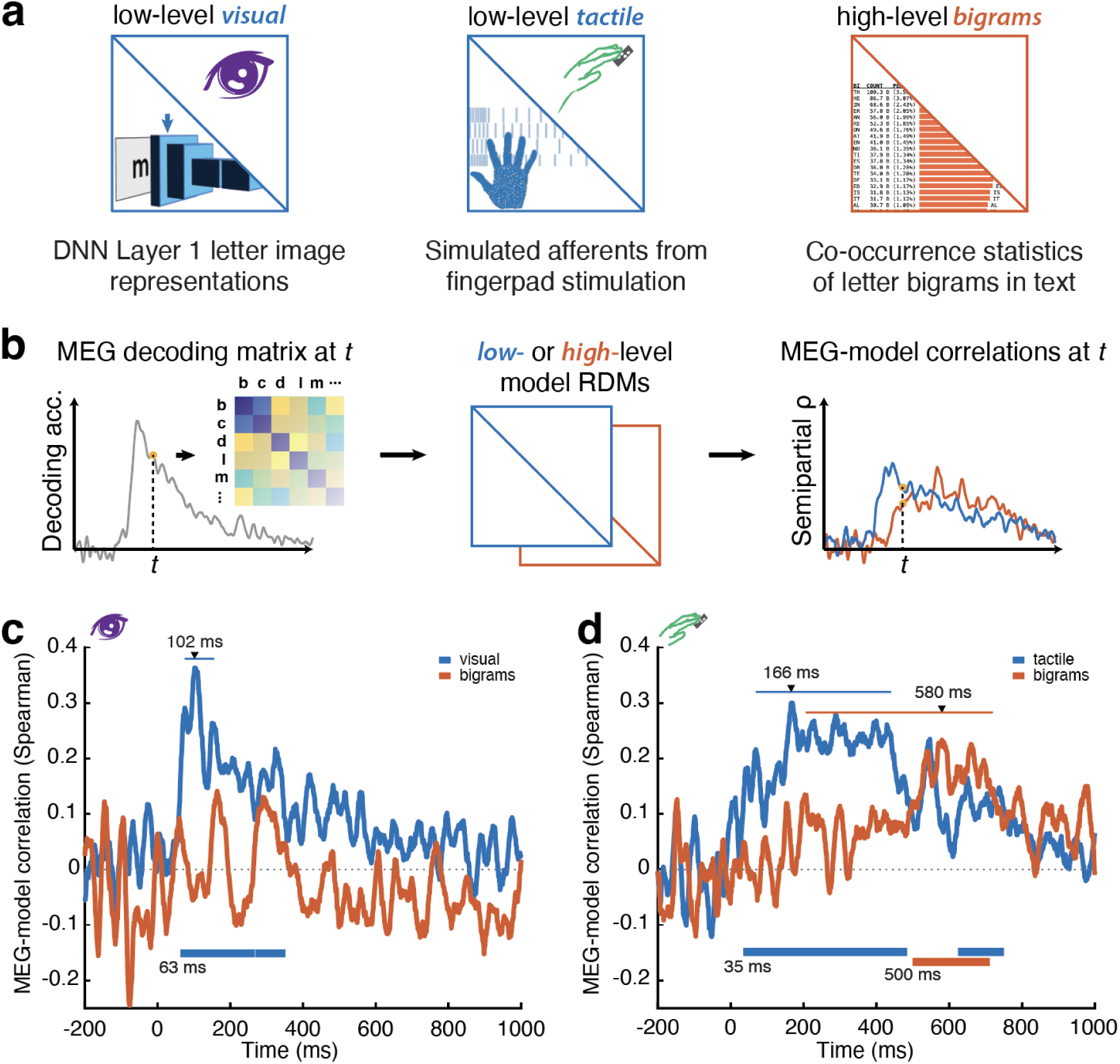
The evolving format of letter representations. (**a**) Models used to test hypotheses about representational format. Low-level (*visual*, *tactile*) models were tested specific to group/modality; high-level *bigram* model capturing text embeddings was tested for both groups. (**b**) Schematic of model comparison between MEG data and models. Comparisons are performed using partial rank-order (Spearman) correlation. (**c,d**) Whole-brain model comparison time courses for sighted and blind participants. Significance testing, significant time point clusters, peaks, and 95% CIs as in Fig. 1. Significance onset times for each model shown at bottom of each plot.

As expected, in the sighted group (Fig. 2c), visual letter representations exhibited a strong and early correspondence with the visual low-level model (Cichy et al., 2016a), beginning at 63 ms (38–132 ms) and peaking at 102 ms (75–155 ms) relative to stimulus onset. However, we did not find significant evidence for correspondence to the high-level bigram model. This pattern of results confirms the contribution of low-level feature representations to individual visual letter processing, while remaining silent about the format of high-level visual letter representations. In contrast, in the blind group (Fig. 2d), both the low-level tactile model and the high-level bigram model corresponded significantly with braille-elicited brain responses. Importantly, the bigram model correlation emerged significantly later than the tactile model correlation for both onset (414 ms; p=0.03) and peak latencies (465; p=0.0046); see Supplementary Table 2 for details. These results characterize the evolving format of letter neural representations in blind brains, revealing a transformation from a low-level tactile to a high-level linguistic format.

### The spatial distribution of letter representations

We next aimed to refine the account of letter representations in sighted and blind brains by assessing their spatial distribution. For this, we decoded source-localized MEG activity in three key regions of interest (ROIs) relevant to visual and braille letter perception (Fig. 3a): 1) Sensorimotor Cortex (*SM*), the cortical point of entry for tactile stimulation (Hamilton and Pascual-Leone, 1998; de Haan and Dijkerman, 2020); 2) Early Visual Cortex (*EVC*), the cortical point of entry for visual stimulation (Hubel and Wiesel, 1962; Felleman and Van Essen, 1991); and 3) left-lateralized Fusiform/Inferotemporal Cortex (*IT*), a high-level part of the ventral visual stream comprising the VWFA and other letter-sensitive regions (Grainger et al., 2008; Dehaene and Cohen, 2011; Reich et al., 2011; Striem-Amit et al., 2012; Lochy et al., 2018). We conducted the time-resolved MVPA analogous to the whole-brain analysis outlined above, but separately for each ROI.

**Figure 3:**
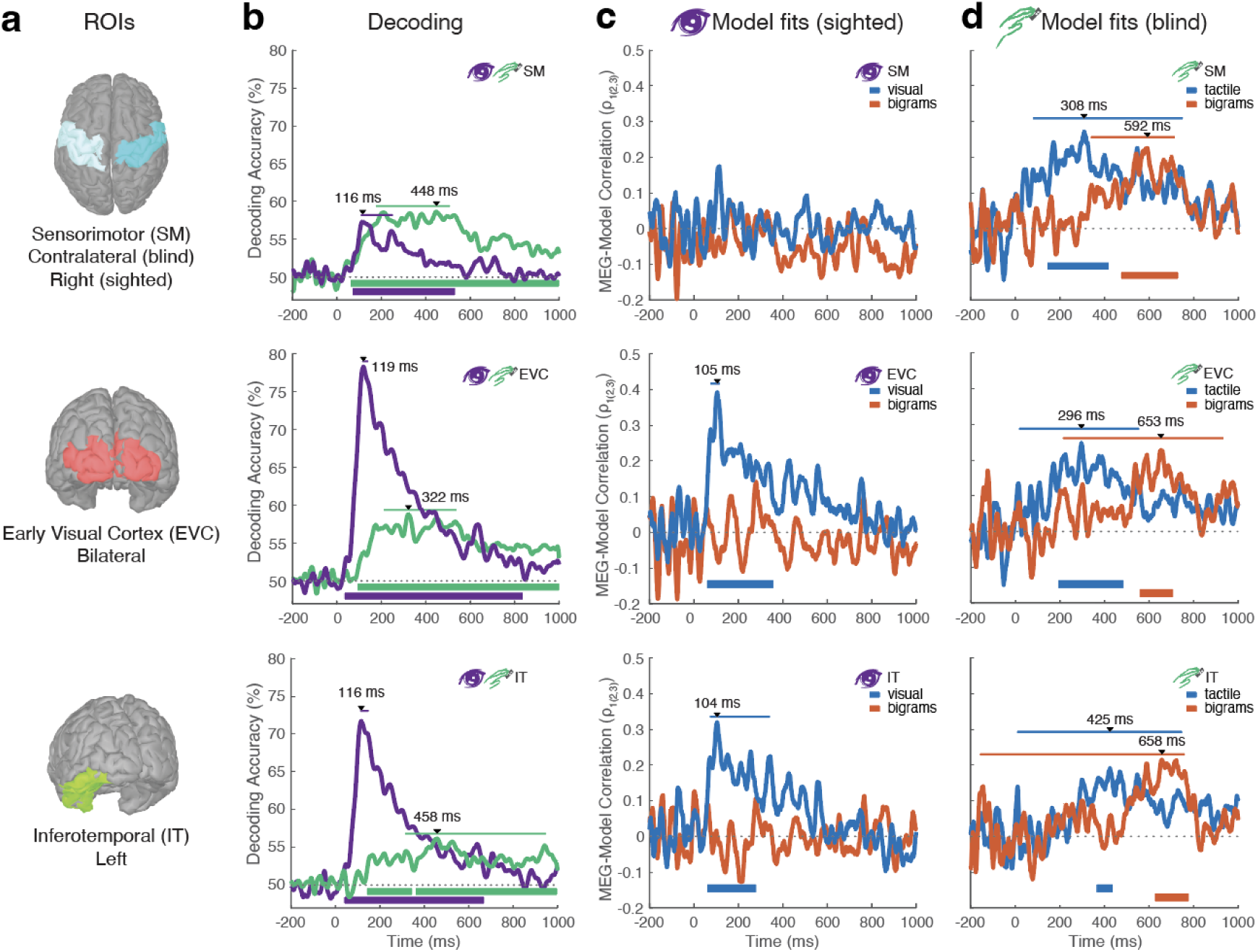
Spatiotemporal distribution of letter representations and formats in key ROIs. (**a**) Key anatomical regions of interest (ROIs) for MEG source localization. *Sensorimotor* (*SM*, *top*) comprises pre- and postcentral gyri on the right (sighted) or stimulation-contralateral (blind) hemisphere. Early visual cortex (*EVC*, *middle*) comprises pericalcarine, lateral occipital, cuneus, and lingual gyrus ROIs. Inferotemporal (*IT*, *bottom*) comprises left-lateralized inferotemporal and fusiform anatomical ROIs. (**b**) ROI-specific MEG decoding of letters color-coded by group, in rows as specified in (**a**). Triangles and thin horizontal bars indicate peaks with 95% CIs; thick bars below curves indicate significant clusters as in previous figures. (**c,d**) ROI-specific MEG-model correlation time courses for group-specific low-level (*visual*, **c**; *tactile*, **d**) and common high-level (*bigrams*) representational models. Peaks, significant clusters, significance onsets, and CIs color-coded by model. Significance was assessed as in whole-brain analyses, via permutation-based cluster-size inference (p < 0.05 cluster-definition threshold, p < 0.05 cluster threshold, one-sided, 500 permutations).

ROI-wise decoding results are shown in Fig. 3b. As expected, in the sighted group we observed strong decoding in EVC and IT, the entrance and late processing stage of visual processing in the brain. We also observe weaker but significant decoding in SM with a similar curve shape as for EVC and IT, likely a result of signal leakage. Comparing decoding onset and peak latencies across ROIs for the sighted group, we found only one significant difference, an earlier onset in EVC than in SM (-37 ms (-3 – 72 ms), p = 0.04; all other p > 0.2).

In contrast, in the blind group, we observed strong and comparable braille decoding effects in SM, EVC and IT, suggesting that all three regions are involved in processing tactile letter representations. We qualitatively observed a systematic increase in onset latency, with shortest latency in SM 63 (28 ms–114 ms), followed by EVC 94 ms (67–149 ms) and finally IT 143 ms (80–399 ms), suggesting a processing cascade. Pairwise comparison of onset latencies across ROIs partially supported this observation, with significant SM–EVC and SM–IT effects (ps<0.05), but not between EVC and IT (p=0.12). Peak latencies did not differ significantly (all p>0.1).

These results clarify the distribution of letter representations in sighted and blind participants, tentatively suggesting a processing cascade for tactile letter representations in the blind from SM over EVC to IT.

### The format of letter representations in SM, EVC and IT

Based on the spatially-resolved MVPA analysis, we next repeated the above analysis linking computational models to neural responses, but separately for each ROI rather than across the whole-brain MEG data as above. In the sighted group (Fig. 3c), for the low-level visual model we observed the expected pattern of results (Grainger et al., 2008; DiCarlo et al., 2012; Cichy et al., 2014): the model predicted activity in EVC and IT as bookends of the feedforward ventral visual stream, but not in sensorimotor cortex (see Supplementary Tables S4 and S5). The high-level bigram model did not predict activity in any ROI. This shows that low-level representations of letters localize to EVC and IT within the regions we examined.

In the blind group (Fig. 3d), both the low-level tactile and the high-level bigram models correlated significantly with MEG decoding patterns in all three ROIs (details see Table S4). Consistent with the outcome of the MVPA analysis, this establishes that all ROIs process braille letters in both low tactile- and high linguistic-level in the blind brain.

Beyond this we detected two further patterns in the results. First, we qualitatively observed that the onset latencies of the correlation time course for the tactile model were earlier than for the bigram model in all three ROIs, consistent with the results pattern observed for the sensor-level analysis above. This observation was statistically ascertained for onset latencies in SM (latency difference 185 ms, p=0.0214), marginally significant in EVC (370 ms, p=0.078), but not supported in IT (p=0.14). This strengthens the view that tactile letter representations transform from a low-level tactile to a high-level linguistic format across time. Second, onset latencies for both models qualitatively followed the same order as for the analysis decoding Braille letters from ROIs: onset latencies were shortest in SM, followed by EVC and then finally IT (respectively: 148, 189, 354 ms for *tactile*; 333, 559, 630 ms for *bigrams*). However, this observation was supported statistically only for the delay between SM and IT (206 ms, p<0.002; see Supplementary Table S5 for details).

Together, our results show that SM, EVC and IT all contribute to processing braille letter representations in the blind brain, and suggests that a spatial hierarchy starting with SM and ending in IT mediates the transformation of earlier low-level tactile into later high-level linguistic representations.

### The inter-region dynamics of letter representations

The above analyses capture representational dynamics for each region separately. However, in a functional network, regions interact and stimulus information is communicated across regions over time, during which we would expect representational formats to be shared between the neural dynamics of the communicating ROIs. Thus, in a final step we interrogated the evolution of letter representations by assessing shared representations across the 3 ROI pairs (i.e. SM and EVC, EVC and IT, and SM and IT) for each group. For this we used a model-based commonality analysis based on variance partitioning (Seibold and McPhee, 1979; Hebart et al., 2018). In brief (Fig. 4a,b), we determined the shared variance between MEG RDMs in a given region pair (e.g. EVC and IT) and either the low- or the high-level model, while discarding the effect of the other model. The analysis yields a commonality coefficient *R*^2^ indicating each model’s unique contribution to the shared variance at all time-point combinations between ROIs, producing a 2D matrix of *R*^2^ values for each model and ROI pair, with respective ROI time points indexing the row and column axes (Fig. 4b). We statistically assessed the *R*^2^ for each model against the other, i.e. the difference score between low- and high-level models and vice versa. The relative onsets (edges) and centroids of the resultant two-dimensional patches of unique model contributions were compared within this space (Fig. 4c; see Methods for details).

**Figure 4.**
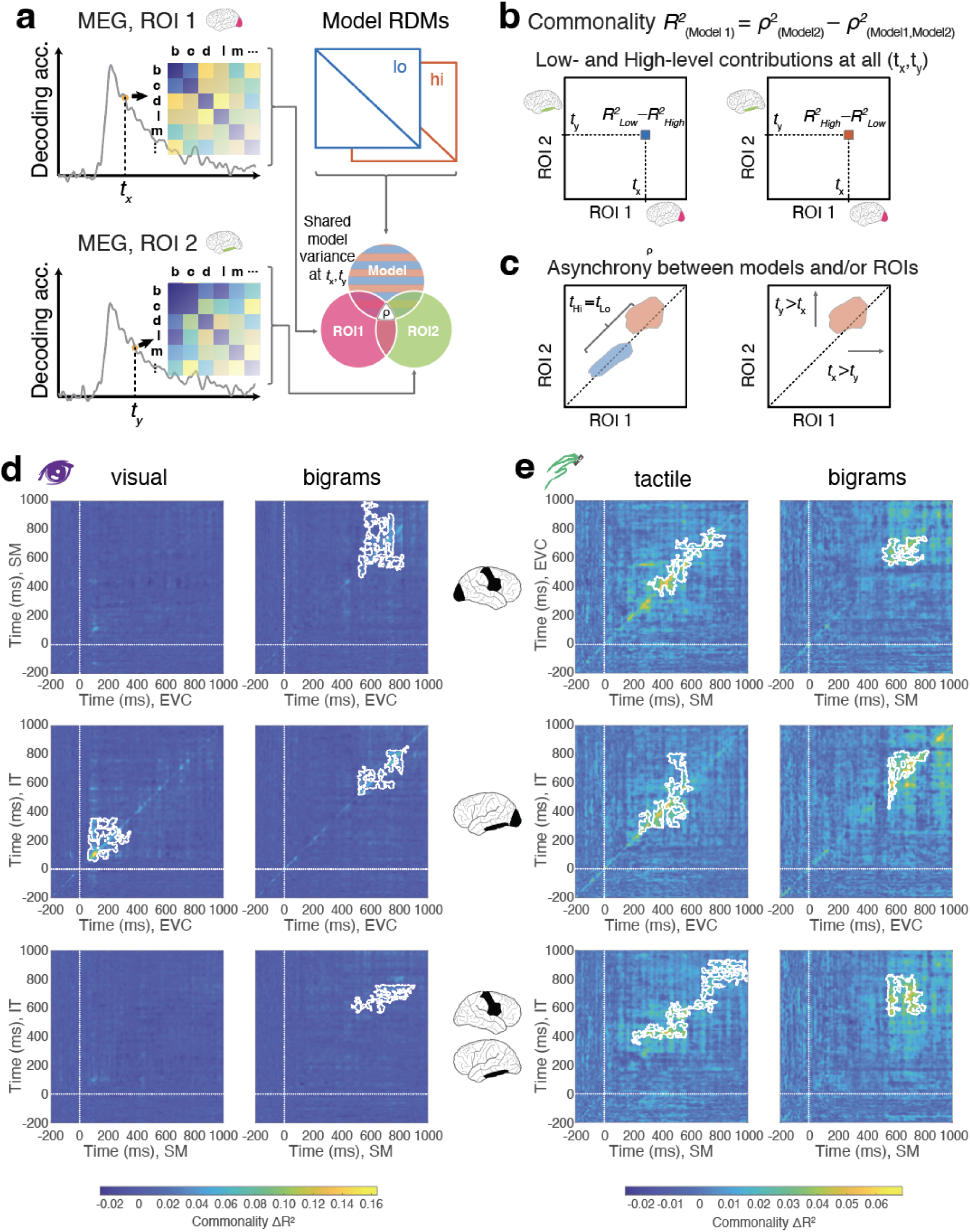
Interregional dynamics of shared letter representations. (**a**) Schematic of commonality analysis identifying shared variance between pairs of ROI-specific MEG data and models. (**b**) Computing the commonality R^2^ for both models across all pairs of time points between MEG ROIs yields a temporal generalization map of uniquely shared variance for each model, expressed as an R^2^ difference score for low (*left*) or high-level (*right*) representations. (**c**) The geometry of the resulting “patches” of significance indicates inter-model differences in temporal dynamics (*left*) or inter-ROI temporal differences (*right*) suggestive of information flow. (**d,e**) Commonality results for sighted and blind groups, respectively, with low- and high-level representations plotted separately for clarity. White dotted lines indicate stimulus onset. White contours outline significant clusters, assessed by 2-d permutation-based cluster-size inference with parameters as in other analyses (p < 0.05 cluster-definition threshold, p < 0.05 cluster threshold, one-sided, 500 permutations).

In the sighted group, we found significant effects for the low-level *visual* model only for EVC and IT (Fig. 4d, left column) between ∼100–400 ms. In contrast, we found significant effects for the high-level *bigram* model for all region pairs emerging (Fig. 4d, right column) between ∼500–900 ms. For the EVC-IT pair, where both models fit the neural data, shared low-level visual model contributions reliably preceded high-level bigram fits in both EVC (onset asynchrony 453 ms, p = 0.002; centroid asynchrony 711 ms, p < 0.002) and IT (onset asynchrony 464 ms, p = 0.044; centroid asynchrony 667 ms, p < 0.002). This suggests that EVC, IT and VWFA partake in representing low-level visual representations, whereas only EVC and IT partake in representing both low-level visual letter representations before high-level bigram representations emerge.

For the blind group (Fig. 4e), we find significant effects for all region pairs, for both the low-level tactile as well as the high-level bigram model. Qualitative inspection suggested that effects for the low-level tactile model emerged prior to the high-level bigram model, but this was not substantiated by statistical analysis (all p > 0.14). Together, the result indicates that EVC, IT and VWFA share low- and high-level braille letter representations.

## DISCUSSION

### Summary

In the present study, we observed the time course of MEG responses to alphabetic braille characters presented to the fingerpads of early-blind participants, and visual printed letters presented foveally to sighted participants. This allowed us to establish the specifics of braille letter representations in blind readers, compared to the visual processing route in sighted subjects. Our main findings are twofold. First, we establish the differential temporal dynamics with which alphabetic letter representations emerge in blind and sighted brains. Second, we detail how letter representations are transformed from a low-level sensory to high-level linguistic format across processing time and cortical regions.

### The temporal dynamics of braille letter representations in blind brains

Previous work characterizing the temporal dynamics of neural response patterns has helped to reveal the functional architecture of visual (DiCarlo and Cox, 2007; Cichy et al., 2014; Yamins and DiCarlo, 2016; Cichy and Kaiser, 2019; Graumann et al., 2022) and auditory (Brodbeck et al., 2018; Ogg et al., 2020; Lowe et al., 2021) processing cascades. Here we thus assessed the temporal dynamics of neural responses to alphabetic responses in braille and print format in blind and sighted brains respectively. Our results revealed different dynamics of letter perception in blind compared to sighted readers, with slower dynamics in the blind group. This difference in processing speed might contribute to the generally slower reading speed for braille compared to print character text (Wetzel and Knowlton, 2000). More generally, our results demonstrate the feasibility of decoding rich single-letter braille representations from MEG, despite suboptimal reading conditions compared to more ecological behavior. A supplementary temporal generalization analysis (King and Dehaene, 2014) further refines this view, showing that both persistent and transient neural representations underlie the observed time courses in both groups and exhibit different dynamics across ROIs (Supplementary Figs. S1, S2). Together these findings invite future research to harvest the power and sensitivity of multivariate methods for research on brain plasticity using braille reading as a model system.

### The format of braille letter representations in blind brains

We found that the brains of blind participants transform representations of braille letters from tactile to linguistic format reflecting the statistics of their pairwise embedding in written text, although each letter was only ever presented as an individual stimulus. This pattern is reminiscent of the gradual abstraction of relevant features from sensory input representations in, e.g., vision (Carlson et al., 2013; Cichy et al., 2014, 2016b) and audition (de Heer et al., 2017; Ogg et al., 2020; Lowe et al., 2021; Giordano et al., 2023). Our results are consistent with previous RSA-based modeling of visual word reading in fMRI (Fischer-Baum et al., 2017), which suggests dissociable visual, orthographic, and semantic processing in ventral occipitotemporal and angular gyrus ROIs. While abstract/semantic representations of single letters are less straightforward than those of whole words (Popham et al., 2021), phrases (Fyshe et al., 2016; Fyshe, 2020), or continuous language (Caucheteux et al., 2023), sublexical processing is tied to visual reading performance in development (Ritchey and Speece, 2006; Acha et al., 2024) as well as adult braille reading (Wilson et al., 2024). Extending earlier work by resolving text-elicited brain responses at the single-letter and millisecond level, we reveal a dynamic and brain-wide network rapidly transforming letter representations through persistent, overlapping representational stages.

Counterintuitively, we found that the *bigram* model contributes to visual print representations in sighted participants, but only in the commonality analysis. Given that the co-occurrence statistics of this model were derived from a visual print corpus, what would explain the relative scarcity of a *bigram*-related signature in sighted versus blind letter responses? The commonality analysis focuses on signals shared across ROI pairs and the model, effectively eliminating non-shared variance. This suggests that aspects of representations reflecting bigram contributions are shared between ROIs, whereas other aspects of the representations are not. These non-shared aspects are arguably strong contributors to representations: for example, coding for low-level visual features in EVC would have been driven by 500 ms of continuous input from a high-contrast uncrowded letter image of typical size for visual neuroscientific studies, but an order of magnitude larger than the 0.2° (or less) subtended by standard printed text (Beier and Oderkerk, 2019). By contrast, braille-reading participants were presented with letters at the same scale they would be typically read. While this design maximized the visual neural signal available for decoding, the proportional imbalance might have caused the visual input to overwhelm higher-level, more abstract signals for individual ROI or whole-brain analyses.

Our approach presents fertile ground for further elucidating the cortical representations activated during braille reading. This could incorporate more and different models, including those derived from behavioral judgments (Haupt et al., 2024) and richer representations elicited by using full words or sentences as stimuli. In addition, while braille students typically learn alphabetic letters first, modern braille text is often rendered in contracted form (D’Andrea et al., 2014); braille-specific models of high-level text representations should account for this when using enriched stimuli. The RSA-based modeling approach could also illuminate evolving representations in braille learners as they improve their proficiency, e.g. by tracking changes in the compositionality of letter combinations (Agrawal et al., 2019, 2020).

### A processing cascade for tactile letter representations in the blind from SM over EVC to IT?

Crossmodal plasticity in early- and congenitally blind braille readers has been a source of debate for decades (Sadato et al., 1996; Büchel, 2003; Amedi et al., 2017; Bedny, 2017; Makin and Krakauer, 2023). Our results inform this debate by suggesting a processing cascade in which tactile braille afferents enter SM and are communicated cortico-cortically to EVC before continuing to IT. This suggests that EVC plays a mediating role between SM and IT, rather than being an epiphenomenal activation. Furthermore, a somatosensory-EVC pathway has been implicated in detecting and identifying tactile/braille stimuli (Hamilton and Pascual-Leone, 1998; Pascual-Leone et al., 1999; Ioannides et al., 2013), but has not been shown directly. Here we provide empirical evidence consistent with this prediction.

While further work is needed to detail the spatiotemporal dynamics of braille reading, our work makes clear predictions about the nature and spatiotemporal profile of braille letter processing that could be experimentally probed with active methods such as TMS (Cohen et al., 1997; Hamilton and Pascual-Leone, 1998). The spatial characterization of these functional networks may also be pursued with fMRI, and link spatially and temporally resolved representations in M/EEG for a spatiotemporally integrated view (Haupt et al., 2024).

### Limitations

Our experimental framework for assessing braille letter perception must be viewed in light of several key limitations in ecological and external validity to real-world context. First, the braille reading conditions encountered by our blind participants were atypical. To precisely control stimulus presentation timing, minimize muscle artifacts, and maximize the signal-to-noise ratio of the MEG recordings via repeated presentations, we presented an alphabetic subset of single letters to our braille readers’ static fingers via refreshable braille display, asking them to omit the active sweeping movements typically observed in braille readers. Still, they identified letters with relative ease (Fig. 1b), suggesting that our results could generalize to braille processing in real-world situations. Second, our setup does not assess higher-level lexical content characteristic of real-world braille text reading as captured in words, sentences and stories. However, the model analyses revealed neural sensitivity to both the letters’ low-level features and their embedding statistics in written text, a convincing signature of literacy even at the single-letter level. Finally, naturalistic reading in proficient braille readers often involves two hands (Martiniello and Wittich, 2022), a cross-hands integration step explored in similar recent single-letter work (Haupt et al., 2024) but not addressed here. Solving the technical challenges of brain measurements in active reading would address all these limitations to reveal the sensorimotor dynamics of continuous text processing.

### Conclusions

In sum, the present study reveals the spatiotemporal and representational dynamics of alphabetic letter processing in blind readers: it involves a transformation from low- to high-level representations that is realized in a cortical network suggestive of a cascade from sensorimotor cortex, over EVC to IT.

## ACKNOWLEDGMENTS

This work was supported by the Vannevar Bush Faculty Fellowship program sponsored by the Basic Research Office of the Assistant Secretary of Defense for Research and Engineering and funded by the Office of Naval Research Grant N00014-16-1-3116 (to A.O.) and the McGovern Institute Neurotechnology Program (to A.O. and D.P.). S.T. was supported by Smith-Kettlewell Institute and the National Eye Institute (Training Grant 5T32EY025201). R.M.C is supported by the Deutsche Forschungsgemeinschaft (DFG; CI241/3-1, CI241/3-3, CI241/3-7 and INST 272/297-1) and by a European Research Council (ERC) starting grant (ERC-2018-STG 803370). We thank K. Ramakrishnan for assistance with early DNN modeling of visual features.

## METHODS

Twelve congenitally or early-blind (7 females; mean age 28.8 y, 5.7 SD) volunteers participated in the study (see Table 1 for details). The maximum age of blindness onset was 3 years. Participants were self-reported fluent daily braille readers and native English speakers. They were either totally blind or had nonspatial light perception; none had had any experience with printed letter reading or visual form generally.. Additionally, 13 sighted volunteers (8 females; mean age 27.1 y, 4.9 SD) with normal or corrected vision were recruited as the visual control group. All participants were compensated for their participation and gave written informed consent in accordance with the guidelines of MIT’s Committee on the Use of Humans as Experimental Subjects.

### Stimuli, task and procedure

Braille stimuli for blind readers comprised 12 single lowercase alphabetic letters (b, c, d, e, l, m, n, o, v, x, y, z), presented via a custom-built single-cell refreshable braille display (model P16; Metec AG, Stuttgart, Germany). The subset was chosen to cover a wide range of letter positions in the alphabet while maximizing stimulus repetitions, and thus signal, within experimental time constraints. During testing, participants rested their index finger on the braille display cell, which presented the stimulus directly to their stationary fingerpads. To reliably peg stimulus onset times to the display’s piezoelectric rods, we affixed a small accelerometer to each participant’s fingernail, registering each impact as a z-axis “spike” synchronized to the MEG recording. Visual stimuli for the sighted group comprised the full alphabet of 26 lowercase alphabetic print letters, rendered in Myriad Pro Bold font using Adobe Illustrator (Adobe, Inc., San Jose, CA) and presented foveally to subtend ∼2° on a rear-projection screen. From this set, the subset of 10 letters corresponding to the braille stimuli was extracted for equivalent analysis to the braille conditions. For both groups, letters were presented in pseudorandom order using custom code and the Psychophysics Toolbox (Brainard, 1997) for Matlab (The MathWorks, Natick, MA). Stimulus duration was 500 ms, with SOA jittered between 1100 and 1300 ms. The experimental sessions comprised approximately 10 runs, resulting in an average of ∼100 presentations per letter condition. Participants performed a vigilance letter-detection task by responding via button press to targets (either an ‘e’ or an ‘o’ for blind, and ‘omega’ (Ω) for sighted) which appeared every 3 to 5 trials. All target trials were excluded from further analyses reported here.

### MEG data acquisition

We scanned participants in an Elekta Neuromag TRIUX MEG scanner (Elekta, Stockholm, Sweden), with continuous whole-brain data acquisition at 1 kHz from 306 sensors (204 planar gradiometers; 102 magnetometers), filtered online between 0.3 and 330 Hz. Head motion was tracked at 330 Hz using five head-position indicator coils affixed to each subject’s scalp, whose location was digitized prior to scanning along with three fiducials and other location markers.

### Data preprocessing

Data were motion-compensated and spatiotemporally filtered offline using Maxfilter software (Elekta, Stockholm, Sweden). All further analysis was conducted using a combination of Brainstorm software (Tadel et al., 2011) and custom analysis scripts, both Matlab-based. From the preprocessed MEG data, we extracted trial epochs for each letter presentation with a prestimulus baseline of 200 ms and 1000 ms post-stimulus onset. For each epoch, the baseline mean was removed and a 30 Hz low-pass filter applied.

### Multivariate analysis of MEG data

We decoded letter identity from trial epochs using a linear support vector machine (SVM; Chang and Lin, 2011 http://www.csie.ntu.edu.tw/~cjlin/libsvm/). For each time point in the epoch, the following analysis was performed: The MEG data yielded a 306-dimensional pattern vector for each of N trials per condition (Fig. 1b). The single-trial pattern vectors were whitened and reduced using PCA to improve SNR, then randomly sub-averaged to yield N_10_=N/10 subaverages per condition, which were then used in a leave-one-out cross-validation approach to train the SVM classifier for every pairwise condition comparison (Cichy et al., 2014; Guggenmos et al., 2018). This process was repeated 100 times, each time with randomized sub-averaging and assignment of training and testing sets. The overall measure of decoding accuracy for a given pairwise comparison and time point was the mean of those 100 permutations. Decoding each pair of conditions (45 pairs for 10 conditions) produced a symmetric 10 x 10 decoding matrix of pairwise decoding accuracies, with the diagonal undefined, for each time point. Thus, analysis of each trial epoch resulted in 1201 such matrices. Interpreting decoding accuracy as a measure of dissimilarity, the decoding matrix is a *representational dissimilarity matrix* (RDM; Kriegeskorte et al., 2008). To generate the grand average decoding time course for the epoch (Fig. 1c), a mean accuracy was computed from the individual pairwise accuracies of the RDM for each time point.

### MEG source estimation and ROI selection

We estimated cortical generators of the MEG signal using Brainstorm’s minimum norm estimate (MNE) method, normalized by noise estimates using a dynamical Statistical Parametric Mapping (dSPM) approach (Dale et al., 2000). For 8 of 12 blind participants, individual T1 structural MRI scans were used to create personalized cortical models; for all remaining subjects, we used Brainstorm’s default cortical model based on the MNI152 template (Fonov et al., 2009). Cortical source estimates were computed on a grid of approximately 15,000 vertices spanning the cortical surface. The forward model was constructed using the overlapping spheres approach (Huang et al., 1999), and MEG signals were mapped onto the cortex using dSPM. From the resulting cortical activity map, we defined ROIs by selecting vertices corresponding to anatomical regions defined in the Desikan-Killiany anatomical atlas (Desikan et al., 2006). For the *Sensorimotor* (SM) ROI, we selected the pre- and postcentral gyrus atlas regions contralateral to the braille-stimulated finger in the blind group (R hemisphere for 9 of 12 subjects); for the sighted group, lacking an analogous selection principle, we used right-hemisphere regions. Our *Early Visual Cortex* (EVC) ROI combines bilateral pericalcarine, lingual, lateral occipital, and cuneus atlas regions. Finally, our *Inferotemporal* (IT) ROI combines left-lateralized inferior temporal and fusiform atlas regions, comprising several left-lateralized regions known to encode higher-level linguistic representations of written text (Reich et al., 2011; Thesen et al., 2012; Fischer-Baum et al., 2017; Haupt et al., 2024). The estimated activity in those regions was subjected to the same sensor-space MVPA decoding analysis described above.

### RDM representational models

To interrogate the representational content of the MEG responses, we computed RDMs corresponding to predicted responses at different levels of analysis. Specifically, we hypothesized sensory-specific low-level response patterns for each group, and a common higher-level abstracted representation for both groups. To model low-level visual representations in the sighted group (“*Visual*” model), we extracted activations elicited by the visual letter images in the first convolutional layer of a deep neural network, AlexNet, pretrained on ImageNet to categorize visual objects (Deng et al., 2009; Krizhevsky et al., 2012; Cichy et al., 2016a). The analogous “*Tactile*” braille model for the blind group computes simulated afferent nerve responses to the spatiotemporal braille indentation pattern on the distal index fingerpad using the Matlab-based touchSIM toolbox (Saal et al., 2017). We set the toolbox stimulation parameters to closely resemble the physical experiment: pins of radius 0.7 mm spaced at 2.5 mm; 0.7 mm indentation in the center of the distal index fingerpad (region D2d) for 500 ms with 20 ms linear ramp; all available innervating fibers (SA1, RA, PC) simulated at full density. From the resulting 945-afferent output, the time-averaged rate vector was extracted to represent the response to each letter. The low-level model RDMs in both cases comprised the pairwise correlation distances (1 – Spearman’s ⍴) between the extracted vectors for each stimulus condition. Finally, because individual letters are semantically impoverished compared to words or longer text, our higher-level model leveraged the statistics of pairwise letter co-occurrences (“*Bigrams*”) over a Google Books English-language text corpus comprising approximately 2.8 trillion exemplars (Norvig, 2013). Bigram co-occurrence frequencies were normalized to the corpus, averaged to be order-independent (e.g. ‘BL’ and ‘LB’ frequencies were averaged), then inverted to index dissimilarity, thus matching the pairwise format of our neural decoding approach. A neural correspondence to the *Bigrams* model would imply high-level sensitivity to letter distributions in written language, rather than to physical letter similarities: more frequent occurrences of a particular bigram reflect lower pairwise dissimilarity in the model and lower MEG decoding accuracy for that letter pair, and vice versa. For all models, Model-MEG correspondence was computed via rank-order semipartial correlation (Spearman’s ⍴) between each computational model RDM and the empirical RDM of the MEG pairwise decoding matrix at each time point, controlling for the effect of the other model (Hebart et al., 2018; Dobs et al., 2019).

### Statistical analysis

The decoding time courses, temporal generalization results, MEG-model correlations, and model-based commonality analyses were tested for significance via permutation tests for cluster-size inference and bootstrap tests for peak and onset time confidence intervals (Nichols and Holmes, 2002; Maris and Oostenveld, 2007; Cichy et al., 2014). The null hypothesis against which correlations or decoding accuracies were tested was 50% (chance) decoding accuracy or 0 correlation between MEG and model RDMs. To generate an empirical null distribution, we created 1000 permutation samples in which each participant’s MEG response was randomly multiplied by +1 or -1, thus allowing the conversion of the data into p-values. We then corrected for multiple comparisons across time points via cluster-size inference (cluster-definition threshold = p < 0.05), using cluster size as the statistic of interest, an analysis more sensitive to temporally extended weak effects than brief strong effects. Clusters were reported as significant if they contained more time points than 95% of the maximal cluster size distribution (cluster-size threshold = p < 0.05). Note that while each individual cluster-based analysis corrects for multiple comparisons across time points, we did not correct the inter-ROI, inter-group, or inter-model analyses for multiple comparisons. Finally, for peak latency and cluster onset distributions, we bootstrapped the participant sample (with replacement) 500 times, repeating the above analysis for each iteration. This produced an empirical distribution of onsets and peak/centroid latencies from which we estimated 95% confidence intervals. For our directional hypotheses (e.g. low-to-high-level representation onset asynchrony between two ROIs), we applied 1-tailed tests.

