## Supplemental Data and Tables for "Touch to text: Spatiotemporal evolution of braille letter representations in blind readers"

#### WHOLE-BRAIN DECODING AND MODEL FIT STATISTICS PER GROUP

##### Table S1: Whole-Brain Decoding Statistics

| **a)** | **Peak latency, ms** | **Onset latency, ms** |
|---|---|---|
| **Visual/sighted** | 120<br>(113–127) | 32<br>(22–75) |
| **Blind/braille** | 329<br>(160–460) | 82<br>(50–125) |
| **b)** | **Peak latency difference, ms** | **Onset latency difference, ms** |
| **Blind minus sighted** | 209<br>(54–390) | 50<br>(-9–93) |
| **P-value** | < 0.002 | 0.078 |

**Table S1: Statistics of time-resolved decoding accuracy analyses. a**) Peak and onset latencies for decoding accuracy curves, with 95% confidence intervals in brackets. **b**) Peak and onset latency differences, with 95% confidence intervals in brackets and p-values computed via bootstrapping.

##### Table S2: Whole-Brain RSA Model Fits

| | | **Peak latency, ms** | **Onset latency, ms** |
|---|---|---|---|
| **Sighted** | **Visual (Low)** | 102<br>(75–155) | 63<br>(38–132) |
| | **Bigram (High)** | N/A | N/A |
| | **High–Low diff, ms** | N/A | N/A |
| | **P-value** | N/A | N/A |
| **Blind** | **Tactile (Low)** | 102<br>(75–155) | 35<br>(-2–273) |
| | **Bigram (High)** | 580<br>(207–720) | 500<br>(178–630) |
| | **High–Low diff, ms** | 465 (48–584) | 414<br>(115–591) |
| | **P-value** | 0.0046 | 0.03 |

**Supplementary Table 2: RSA model fit statistics for each model and model comparison.** RSA model comparison peak and onset latencies are displayed by model and group. Bootstrapped 95% confidence intervals in brackets and p-values as in previous figures and tables.

#### ROI DECODING AND MODEL FIT STATISTICS PER GROUP

##### Table S3: ROI-Specific Decoding Timecourse Statistics

| a) | ROI | Peak latency, ms | | Onset latency, ms | |
|---|---|---|---|---|---|
| Sighted | EVC | 119 (114–140) | | 35 (21–66) | |
| | SM | 116 (108–251) | | 72 (27–102) | |
| | IT | 116 (112–149) | | 40 (15–73) | |
| Blind | EVC | 322 (212–537) | | 94 (67–149) | |
| | SM | 448 (176–508) | | 63 (28–114) | |
| | IT | 458 (315–947) | | 143 (80–399) | |
| **b)** | **ROI comparison** | **Peak latency difference, ms** | **P-value** | **Onset latency difference, ms** | **P-value** |
| Sighted | EVC – SM | -3 (-10–128) | 0.776 | -37 (-3–72) | 0.04 |
| | IT – EVC | -3 (-12–37) | 0.226 | 5 (-12–37) | 0.226 |
| | SM – IT | 0 (-131–32) | 0.706 | -32 (-74–16) | 0.922 |
| Blind | EVC – SM | 126 (-183–336) | 0.454 | 31 (-112–4) | 0.046 |
| | IT – EVC | 136 (-75–630) | 0.234 | 80 (-9–296) | 0.124 |
| | SM – IT | 10 (-14–749) | 0.136 | 94 (-1–334) | 0.028 |

**Supplementary Table 3: Statistics of ROI-specific time-resolved decoding accuracy analyses. a**) Peak and onset latencies for ROI-specific decoding accuracy curves. **b**) Between-ROI peak and onset latency differences, with sign in the direction of the first ROI in the comparison label (e.g., positive latency difference for "EVC–SM" means EVC has later peak/onset). 95% confidence intervals (in parentheses) and significance computed by bootstrapping participant sample, 500 iterations, 1-tailed test.

**Table S4: RSA Model Fit Statistics**

| | a) | Model | | b) Onset difference, ms | |
|---|---|---|---|---|---|
| **Sighted** | **Region** | **Visual** | **Bigram** | **Bigram minus visual** | **P-value** |
| | SM | N/A | N/A | N/A | N/A |
| | EVC | 60 (40–93) | N/A | N/A | N/A |
| | IT | 60 (45–131) | N/A | N/A | N/A |
| **Blind** | **Region** | **Tactile** | **Bigram** | **Bigram minus tactile** | **P-value** |
| | SM | 148 (2–285) | 333 (286–509) | 185 (119–453) | 0.0214 |
| | EVC | 189 (100– 650) | 559 (177–723) | 370 (-58–472) | 0.0778 |
| | IT | 354 (241–705) | 630 (176–647) | 276 (-218–384) | 0.1399 |

**Supplementary Table 4: Statistics of the RSA model fit analysis for each model and model comparison.** **a**) Onset latencies for RSA results for the two different models (visual & bigram), for the sighted and blind participants, for the cortical regions Sm, EVC and IT (with 95% confidence intervals in brackets) as reported in Fig. 3c-d. **b**) Onset latency differences in model fits (with 95% confidence intervals in brackets) and p-values (stats details).

**Table S5: Inter-Region RSA Model Fit Onset Latency Statistics**

| a) Sighted | | Model | | | |
|---|---|---|---|---|---|
| | | Visual | | Bigram | |
| | Region Comparison | Onset latency difference | P-Value | Onset latency difference | P-Value |
| | EVC – SM | N/A | N/A | N/A | N/A |
| | IT – EVC | 0<br>(-75–14) | 0.250 | N/A | N/A |
| | SM – IT | N/A | N/A | N/A | N/A |
| b) Blind | | | | | |
| | | Tactile | | Bigram | |
| | Region Comparison | Onset latency difference | P-Value | Onset latency difference | P-Value |
| | EVC – SM | 41<br>(-86–494) | 0.1889 | 226 | 0.1093 |
| | IT – EVC | 165<br>(-197–513) | 0.1042 | 71 | 0.2781 |
| | SM – IT | 206<br>(10–612) | <0.002 | 297 | 0.1155 |

**Supplementary Table 5: Statistics of the RSA model fit analysis comparing onset latencies between regions. a**) Onset latencies for RSA correlation curves for the two different models (visual & bigram), for the sighted and blind participants, comparing the three regions SM, EVC and IT (with 95% confidence intervals in brackets).

#### INTER-ROI COMMONALITY ANALYSES PER GROUP AND ROI PAIR

##### Table S6: Inter-ROI Commonality Analysis Statistics

| | Low-level (tactile/visual) | | | | High-level (bigrams) | | | |
|---|---|---|---|---|---|---|---|---|
| **a) Sighted** | **Onset X** | **Onset Y** | **Centroid X** | **Centroid Y** | **Onset X** | **Onset Y** | **Centroid X** | **Centroid Y** |
| **EVC→IT** | 57 | 53 | 166 | 207 | 510 | 517 | 877 | 874 |
| | [20 132] | [19 73] | [134 571] | [179 561] | [182 707] | [-89 703] | [562 782] | [445 798] |
| **EVC→SM** | N/A | N/A | N/A | N/A | 516 | 466 | 656 | 720 |
| | N/A | N/A | N/A | N/A | [33 651] | [-51 663] | [530 779] | [543 818] |
| **SM→IT** | N/A | N/A | N/A | N/A | 466 | 565 | 702 | 683 |
| | N/A | N/A | N/A | N/A | [124 633] | [-110 687] | [436 774] | [486 831] |
| **b) Blind** | | | | | | | | |
| **SM→EVC** | 327 | 336 | 560 | 571 | 475 | 548 | 637 | 687 |
| | [2 629] | [-67 635] | [303 805] | [315 751] | [142 631] | [-43 634] | [361 782] | [359 792] |
| **EVC→IT** | 242 | 241 | 449 | 480 | 546 | 417 | 649 | 699 |
| | [121 490] | [158 526] | [310 698] | [325 726] | [-73 804] | [-8 623] | [114 915] | [356 745] |
| **SM→IT** | 192 | 342 | 645 | 655 | 538 | 540 | 658 | 691 |
| | [0 684] | [159 673] | [363 855] | [397 831] | [130 669] | [22 627] | [520 799] | [477 758] |

**Supplementary Table 6: Commonality analysis statistics by ROI pair.** Onsets and centroids of significant clusters between low- and high-level models for each ROI comparison in Sighted (**a**) and Blind (**b**) groups. Confidence intervals computed as in other analyses, via bootstrapping (500 iterations). X and Y indicate the X- and Y-axes of the time generalization plots respectively, and thus denote the respective ROIs in the order given in the leftmost column. For clarity, color codes indicate groups (purple = Sighted; green = Blind) and models (blue = low-level; orange = high-level).

#### Table S7: Between-Model Commonality Statistics per ROI Comparison

| | High-Low Model Differences (Within ROI Pair Comparison) | | | | | |
|---|---|---|---|---|---|---|
| **a) Sighted** | **Onset X** | **Onset Y** | **P-value** | **Centroid X** | **Centroid Y** | **P-value** |
| **EVC→IT** | 453 | 464 | 0.002, 0.044 | 711 | 667 | <.002, <.002 |
| | [159 650] | [-129 632] | | [104 588] | [112 563] | |
| **EVC→SM** | N/A | N/A | N/A | N/A | N/A | N/A |
| | N/A | N/A | N/A | N/A | N/A | N/A |
| **SM→IT** | N/A | N/A | N/A | N/A | N/A | N/A |
| | N/A | N/A | N/A | N/A | N/A | N/A |
| **b) Blind** | | | | | | |
| **SM→EVC** | 148 | 212 | 0.142, 0.221 | 77 | 116 | 0.201, 0.183 |
| | [-274 472] | [-278 499] | | [-217 371] | [-144 357] | |
| **EVC→IT** | 304 | 176 | 0.331, 0.506 | 200 | 219 | 0.153, 0.375 |
| | [-366 541] | [-427 386] | | [-310 428] | [-266 310] | |
| **SM→IT** | 346 | 198 | 0.194, 0.36 | 13 | 36 | 0.283, 0.417 |
| | [-326 554] | [-413 383] | | [-238 338] | [-247 284] | |

**Supplementary Table 7: Between-model commonality statistics by ROI pair.** Relative onsets and centroids of significant clusters between models for each ROI pair in Sighted (**a**) and Blind (**b**) groups. X and Y indicate the X- and Y-axes of the time generalization plots respectively, and thus denote the respective ROIs in the order given in the leftmost column. Color code indicates groups (purple = Sighted; green = Blind).

#### Table S8: Within-Model Commonality Comparison per ROI Comparison

| | Diff ROI Timing (Within Model) | | | | | | | |
|---|---|---|---|---|---|---|---|---|
| | Low-level (tactile/visual) | | | | High-level (bigrams) | | | |
| **a) Sighted** | **Onset Y-X** | **P-value** | **Centroid Y-X** | **P-value** | **Onset Y-X** | **P-value** | **Centroid Y-X** | **P-value** |
| **EVC→IT** | -4 | 0.432 | 41 | 0.352 | 7 | 0.584 | -3 | 0.42 |
| | [-61 28] | | [-102 134] | | [-482 103] | | [-160 147] | |
| **EVC→SM** | N/A | N/A | N/A | N/A | -50 | 0.752 | 64 | 0.336 |
| | N/A | N/A | N/A | N/A | [-337 252] | | [-90 165] | |
| **SM→IT** | N/A | N/A | N/A | N/A | 99 | 0.244 | -19 | 0.272 |
| | N/A | N/A | N/A | N/A | [-459 442] | 0.762 | [-151 339] | 0.728 |
| **b) Blind** | | | | | | | | |
| **SM→EVC** | 9 | 0.326 | 11 | 0.323 | 73 | 0.342 | 50 | 0.277 |
| | [-218 300] | | [-120 205] | | [-308 313] | | [-81 285] | |
| **EVC→IT** | -1 | 0.345 | 31 | 0.256 | -129 | 0.653 | 50 | 0.653 |
| | [-190 314] | | [-76 213] | | [-529 347] | | [-356 298] | |
| **SM→IT** | 150 | 0.3 | 10 | 0.345 | 2 | 0.67 | 33 | 0.534 |
| | [-245 361] | | [-117 183] | | [-394 239] | | [-171 174] | |

**Supplementary Table 8: Within-model commonality statistics by ROI pair.** Onsets and centroids relative to diagonal for low- and high-level model representations in Sighted (**a**) and Blind (**b**) groups. X and Y indicate the X- and Y-axes of the time generalization plots respectively, and thus denote the respective ROIs in the order given in the leftmost column. Color codes indicate groups (purple = Sighted; green = Blind) and models (blue = low-level; orange = high-level).

### TEMPORAL GENERALIZATION DECODING AND MODEL FITS

#### Temporal Generalization Decoding

The grand average decoding accuracy time courses shown in Main Figs. 1 and 3 suggest both dynamically changing and persistent representations: they include rapid changes such as the initial steep rise, as well as periods of apparent stability such as the "plateau" in braille letter decoding at ~150–600 ms. However, aggregate decoding accuracy does not index the underlying response patterns; thus, to more concretely interrogate the dynamics of letter-evoked brain responses, we applied temporal generalization (TG) analysis (King and Dehaene 2014; Cichy, Pantazis, and Oliva 2014), an extension of time-resolved MVPA in which classifiers are trained and tested on all combinations of time-points in the epoch, rather than on the same time points as in Main Fig. 1d. The rationale is that for persistent representations, a classifier trained at one time point will succeed across many different time points; for dynamic representations, training and testing will only succeed on the same or very similar time points. Across all pairwise combinations, temporal generalization yields two-dimensional matrices of classification accuracy indexed in rows and columns by the time points used for training and testing (see Fig. S1a below). In sighted subjects (Fig. S1d, below), TG dynamics vary with ROI, with a weaker signal in SM compared to the other two, but are consistent with the combination of transient (on-diagonal) and sustained (off-diagonal) elements reported in previous visual temporal generalization results (Isik et al. 2014; Cichy, Pantazis, and Oliva 2014; King and Dehaene 2014). In blind participants (Fig. S2a), decoding accuracy remains highest along the diagonal across the whole time course up to at least 500 ms, negating the idea of the flat braille decoding plateau from ~150–600 ms as observed in Fig. S2a reflecting fully persistent representations. We also observe no cardinal (horizontal/vertically extended) bands of significance as for sighted letter processing, suggesting that the low-level response tied to stimulus duration is more transient or creates a fainter MEG signal compared to its visual counterpart. However, we observe progressively more persistent dynamics across time, as indexed by the broadening of significant classification across time, i.e. increasing decoding accuracy away from the diagonal, with strong persistence in the late time course (Fig. S2a). Statistical significance was assessed as with main text analyses, via permutation-based cluster-size inference ($p < 0.05$ cluster-definition threshold, $p < 0.05$ cluster threshold, one-sided, 500 permutations), with significant clusters outlined in the TG maps below.

**TG Model Fitting**

As with the 1-d time-resolved grand average decoding curves, each time point of the TG matrix represents an averaged RDM of pairwise decoding accuracies. Thus, we interrogated their content in a similar fashion, via correlation with our low- and high-level model RDMs (Main Fig. 2, Fig. S1b-c below). For the sighted group, as with the 1-d results, only EVC and IT ROIs correlated significantly with the *visual* model, and none correlated significantly with the *bigrams* model (Fig. S1e,f). Note that clusters of significant *visual* model correlation are more elaborated in EVC, including a diagonally oriented portion and cardinal bands consistent with stimulus duration; while significant IT correlation appears more temporally punctate and, qualitatively, onsets later compared to the EVC pattern. In the blind group, both the *tactile* and *bigrams* models correlated significantly with TG decoding matrices for each ROI (Fig. S2b,c). Notably, the *tactile* correlation appears to undergo substantive transformation from SM to EVC to IT, with progressively later significance onset and progressively more sustained dynamics in IT compared to the tightly diagonal-bound correlation cluster in SM. EVC displayed elements of both. The *bigrams* model consistently emerged after ~300 ms in all ROIs, with a broadly consistent "width" (off-diagonal persistence) of 300–400 ms.

Overall, the TG decoding and modeling results corroborate the 1-d time-resolved analysis and further suggest evolving dynamics of pattern representations over the course of the hypothesized pathway from early sensory cortex to IT. The results also indicate simultaneously active low- and high-level letter representations in the ROIs, suggesting a temporally multiplexed processing cascade not strictly following a monotonic feedforward scheme.

#### Figure S1: ROI Decoding and Model Fits for Sighted Group

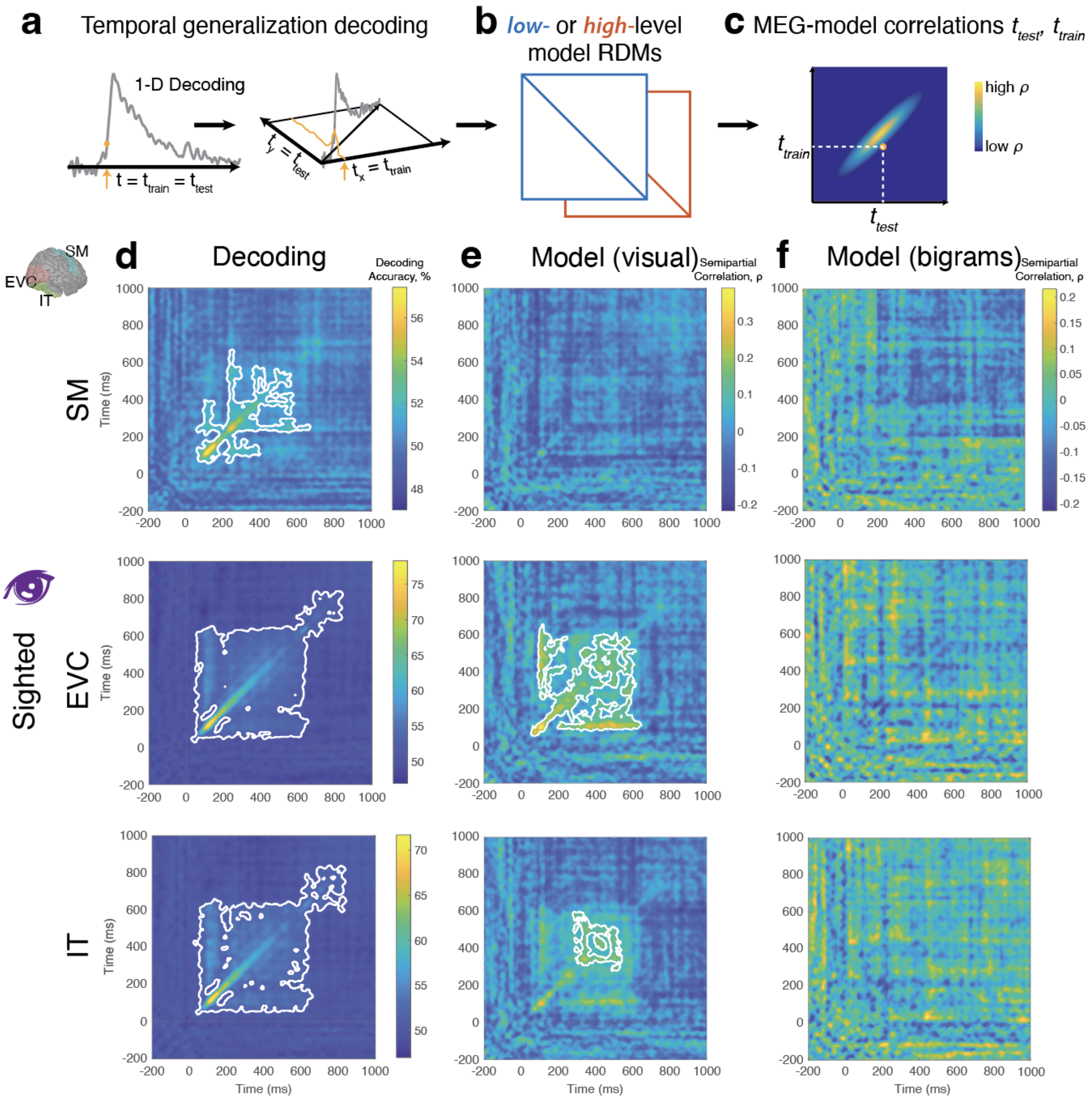

**Supplementary Figure S1: ROI-wise temporal generalization decoding for Sighted group.** (**a**) Overview of temporal generalization (TG), in which classifiers are trained and tested across all time points in the epoch. The resulting TG matrix comprises MEG decoding RDMs, as with the 1-d case in Main Fig. 2, and can be compared with models (**b**) to yield 2-d MEG-model correlation matrices (**c**). (**d**) TG decoding accuracy for each ROI, alongside corresponding low-level visual (**e**) and high-level bigram (**f**) model fits. Model correlation colorbars apply to whole column. White contours outline significant decoding or correlation clusters. Significance assessed as in main text, via permutation-based cluster-size inference ($p < 0.05$ cluster-definition threshold, $p < 0.05$ cluster threshold, one-sided, 500 permutations).

#### Figure S2: ROI Decoding and Model Fits for Blind Group

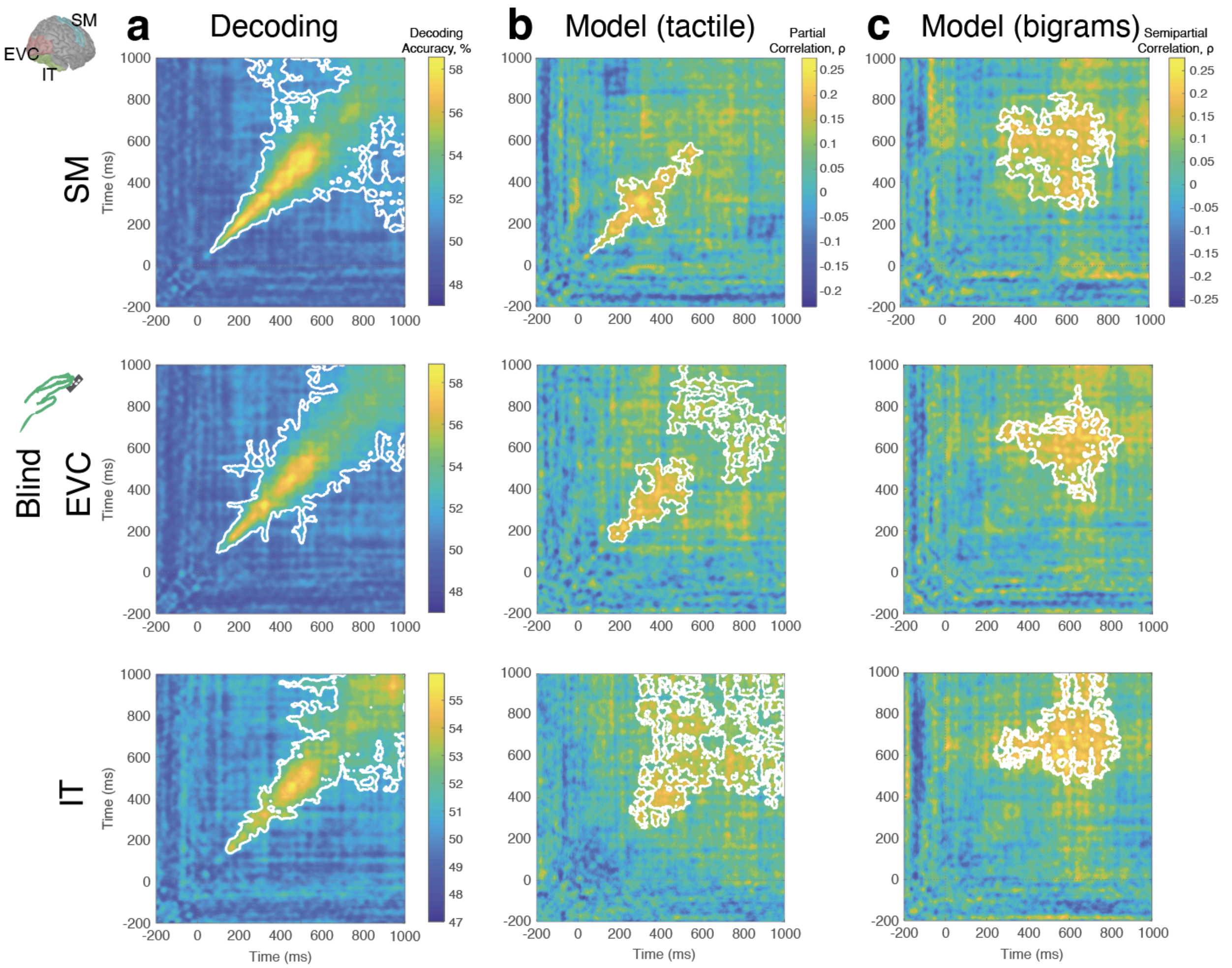

**Supplementary Figure S2: ROI-wise temporal generalization decoding for Blind group.** (**a**) Temporal generalization decoding accuracy for each ROI, alongside corresponding low-level tactile (**b**) and high-level bigram (**c**) model fits. Model correlation colorbars apply to whole column. White contours outline significant decoding or correlation clusters. Significance assessed as in main text, via permutation-based cluster-size inference ($p < 0.05$ cluster-definition threshold, $p < 0.05$ cluster threshold, one-sided, 500 permutations).
